# Field-testing olfactory ability to understand human olfactory ecology

**DOI:** 10.1101/270744

**Authors:** Kara C. Hoover, Denisa Botescu, Piotr Fedurek, Sophie Aarts, J. Colette Berbesque

**Affiliations:** Department of Anthropology, University of Alaska, Fairbanks, USA; Department of Anthropology, University College of London, London WC1H 0BW; Centre for Research in Evolutionary, Social and Inter-Disciplinary Anthropology, University of Roehampton, London SW15 4JD U.K.

**Keywords:** Smelling in the wild, sex differences in olfactory ability, odor identification, olfactory ecology, functional olfaction

## Abstract

**Objectives:** We know little about human olfactory ability in natural settings because current knowledge derives from lab-based studies using non-representative samples of convenience. The primary objective was to use a validated lab tool, the five-item odor identification test, to assess variation in olfactory ability in different environments.

**Methods:** Using the five-item test, we conducted two repeated measures experiments that assessed participant ability to correctly identify an odor source in different odor environments. We also examined consistency in odor labelling due to documented potential bias from idiosyncrasies in odor terms.

**Results:** We found no variation in olfactory ability due to environment, but this may be due to methodological biases. First, subjective bias results from idiosyncratic differences in participant labelling and researcher coding of answer correctness. Second, better ability to learn odors may provide an advantage to women. Third, reducing positive female learning bias by analyzing consistency in response (regardless of correct odor source identification) fails to assess functional olfactory ability. Functional olfactory ability (naming correct odor source) is significantly better in females, especially in food-rich odor environments.

**Conclusions:** Environment was not a significant factor in olfactory ability in this study but that result may be confounded by methodological biases. We not recommend odor identification as a field tool. Functional olfactory ability exhibits a sex-based pattern but consistency in recognizing the same odor does not. Food-rich odors may enhance olfactory ability in females. We discuss evolutionary and ecological implications of superior female functional olfactory ability relative to food foraging activity.

## INTRODUCTION

The vast majority of our knowledge on human olfaction derives from lab studies conducted in odor-controlled environments using samples of convenience, typically white educated industrialized rich and democratic or WEIRD populations (Henrich et al., 2010)— often students (Hanel et al., 2016). While ethnographic work in anthropology has examined olfaction, the focus has been on odor symbolism rather than olfactory ability or olfactory biology (for an overview, see Classen, 1993; Classsen et al., 1994). Meanwhile, the vast majority of human olfactory experience is in dynamic non-western and non-industrialized odorscapes. The limited research done on olfactory ability in different ecological settings suggests that non-industrial populations may have superior olfactory sensitivity but not discrimination abilities (Sorokowska et al., 2014; Sorokowska et al., 2013). While these studies provide preliminary insights into what natural human variation in olfactory ability might be like, comparing data collected in the field to those in the lab challenges interpretation given the unknown variables in the field influencing test outcomes. Thus, we know little about how humans detect, discriminate, and identify odors in mixtures in natural settings—for humans, natural settings include the built environment, both urban and rural.

Lab data provide insights into smelling odor mixtures. First, while we may detect many to most odors in a complex odor mixture, the whole of that information is not transduced to the brain due to the inhibition of some olfactory receptor cells via receptor competition (Jinks et al., 1999). Even if all odors in the mix are subliminally processed in the cerebrum, only some are perceived (T. Hummel et al., 2013). Second, there may be an upper limit to the number of odors detectable individually from mixes—one experiment suggested that more than 30 equal-intensity odorants are perceived as an ‘olfactory white’ (Weiss et al., 2012). Third, the ability to detect single odors in mixtures may be improved by experience (C. Sinding et al., 2015) and training (Barkat et al., 2012; Morquecho-Campos et al., 2019). Fourth, there may be ecological pressures that have acted on the filtering mechanisms of the olfactory system. There is a neuronal pattern that suggests we are particularly tuned to food odors (Saraiva et al., 2019). Thus, the olfactory system is limited by its ability to integrate complex whole mixtures into perception and driven by attention to specific compounds. Perhaps environmental variation outside the lab is not a disruptive confounding factor. By widening data collection to field-based settings, a representative sample would be possible and allow us to understand the extent of natural human variation in olfactory ability.

The principal challenge in generating ecologically salient data is the lack of field-based methods (Kern et al., 2014). This problem with modifying a lab-based test for deployment in the field was first noted in the discipline of anthropology during the ethnographic fieldwork in the Torres Straits at the turn of the century (Myers, 1901). The only field-validated method is the Olfactory Function Field Exam (OFEE), an 11-minute olfactory function test that uses a modified and abbreviated version of Sniffin’ Sticks—two odors for threshold detection (Kern et al., 2014) and five odors for identification (Mueller et al., 2006). As part of The National Social Life, Health, and Aging Project in the United States, the OFFE was validated on a cohort of 2,304 older adults and conducted in participants’ homes following a short odor training session. Results were nearly identical to testing conducted in labs on a similarly aged cohort but not using repeated measures on the same subjects (Kern et al., 2014). OFFE validation results suggest that between-subjects environmental variation was not significant. Olfactory adaptation down-regulates receptor stimulation when odors are constant, however, which means that OFFE subjects were not smelling their home environments and the effect from variation would be small (Auffarth, 2013; P. Dalton, 2000).

Our goal was to assess the impact on olfactory ability from a dynamic environment to which subjects were not adapted. We needed a tool that was highly portable and easy to administer in the field. The ease of administering the two most accurate tests, threshold and discrimination, in the field without a table was prohibitive. That left odor identification. There are three validated short tests of olfactory ability that rely solely on odor identification—a five odor Sniffin’ Sticks test (Mueller et al., 2006), a three odor Sniffin’ Sticks test (Thomas Hummel et al., 2010), and a three-odor test using microencapsulation (Jackman et al., 2005). We used the five-item test in two repeated measures studies: a field study comparing olfactory ability in a low odor environment to 1) a polluted environment and 2) a high odor environment; a lab study comparing olfactory ability in a low odor control lab to a high odor experiment condition.

## MATERIALS AND METHODS

### Materials

Our five-item test included four of the original five-item test odors: peppermint, orange, rose, fish (Mueller et al., 2006). Leather was replaced by garlic because it had a lower identification rate in the UK (Neumann et al., 2012). The field study was conducted in August 2018 and the lab study in February 2019. Materials were purchased in July 2018 and stored at the University of Roehampton at a temperature within the recommended manufacturer’s range.

### Screening and Testing Protocols

Testers wore cotton gloves during testing and wore no scented products. Each odor was moved across the nostrils slowly for three seconds. Participants were asked to describe the odor via free response then choose one of 22 terms from a verbal prompt card: five odor terms (one correct per odor), 15 distractor odor terms (none correct for any odor), ‘undefinable’, and ‘no odor’ (see Table 2 of Mueller et al., 2006). The field study consisted of three tests per participant and the lab study consisted of two. Between-test gaps were small (25-30 minutes for field, 15-20 for lab).

**Table 1:**
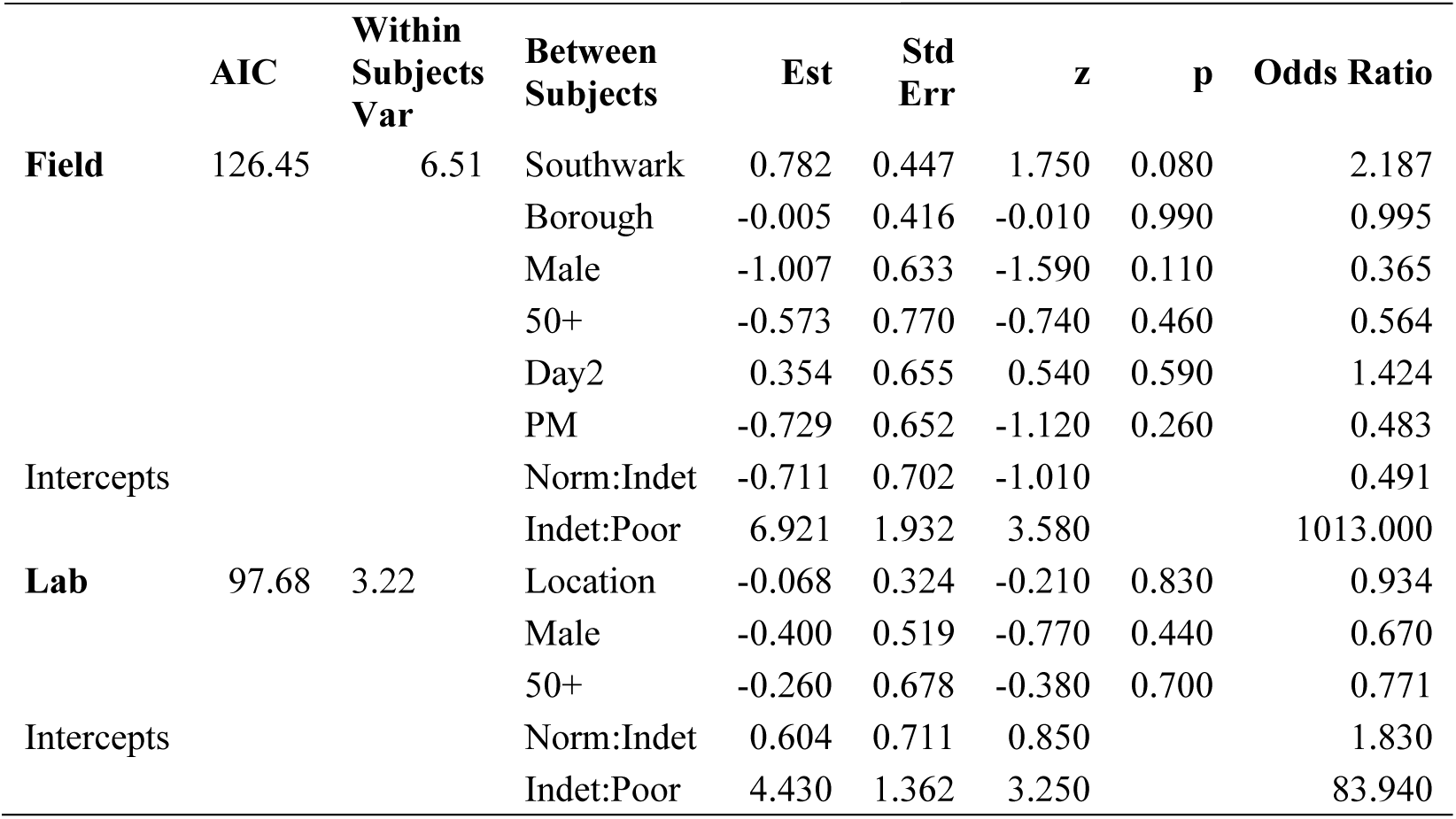
Ordinal regression results.

**Table 2:** Sign Test Results.

| Subset | Contrast | S | p-value | 95%CI |  |
| --- | --- | --- | --- | --- | --- |
|  |  |  |  | Lower | Upper |
| All | Tate:Borough | 3 | 0.100 | 0.857 | 0.951 |
| Female | Tate:Borough | 0 | 0.060 | 0.854 | 0.961 |
| Male | Tate:Borough | 3 | 1.000 | 0.854 | 0.961 |
| All | Tate:Southwark | 3 | 0.500 | 0.901 | 0.957 |
| Female | Tate:Southwark | 0 | 0.100 | 0.922 | 0.973 |
| Male | Tate:Southwark | 3 | 1.000 | 0.820 | 0.961 |
| All | Control:Experimental | 6 | 1.000 | 0.939 | 0.976 |
| Female | Control:Experimental | 5 | 0.700 | 0.939 | 0.976 |
| Male | Control:Experimental | 1 | 1.000 | 0.857 | 0.951 |

### Data Coding and Statistical Analysis

Data are shared at <u>10.6084/m9.figshare.11764272</u> under the CC BY 4.0 license. All data were analyzed and graphics plotted in R for Windows v3.6.1 (Charif et al., 2007) using the R Studio GUI v1.2.1335 (RStudio Team, 2015). Responses were coded as correct, near miss (a related item like lime for orange), far miss (a more general term like citrus for orange), and incorrect (an unrelated item like raspberry for orange) following Cain (1979). Between-coder agreement (JCB, KCH) was tested by Cohen’s kappa (κ) using *irr* (Gamer et al., 2012) in the lpsolve package (Berkelaar, 2015). The total correct provided an ordinal level variable describing olfactory ability (Mueller et al., 2006):

- Poor (0): severely impaired or no sense of smell
- Indeterminate (1 to 3): further testing required
- Normal (4-5 correct): normal or slightly reduced sense of smell

The effect on olfactory ability from differing environments was estimated using ordinal regression (AKA logistic regression and cumulative link models), specifically, *clmm* in the ordinal package (Christensen, 2019). Logistic regression offers a powerful alternative to traditional ordinal tests (e.g., Mann-Whitney U, Wilcoxon signed rank sums, Kruskal Wallis) because it allows the estimation of fixed effects on the dependent variable and estimation of repeated measures via a random term (McCullagh, 1980)—an ordinal version of mixed-model linear regression or repeated measures ANOVA. We used the proportional odds model which assesses the probability of being in *x* ranked category or lower versus being in any higher ranked category with olfactory rank as the thresholds between categories. We included additional fixed effects—sex because females perform better in lab studies (Oliveira-Pinto et al., 2014; Sorokowski et al., 2019) and age because olfactory ability declines after age 50 in lab studies (Martinez et al., 2017; Charlotte Sinding et al., 2014).The field study had two additional effects, day and time of testing, due to potential temporal variation in test environments. The proportional odds effects plots were generated in the effects package (Fox et al., 2009) using output from *polr* in MASS (Venables et al., 2002).

Odor identification is the hardest Sniffin’ Sticks task due to challenges in odor term recall and lack of social consensus in odor naming; for example, the association of cloves with the dentist or lemon and rose with cleaning supplies may result in marking an identification incorrect even though the participant has actually identified the odor but not used the veridical (or true) label, e.g., orange for orange rather than citrus or fruity (Cornell Kärnekull et al., 2015; Jönsson et al., 2016). Thus, we also analyzed the data for internal consistency in odor term (whether correct, near miss, far miss, incorrect) via an ordinal ranked variable where a rank of 0 (inconsistent terms across trials) or 1 (consistent terms across trials). If an individual could not smell the odor in any trial, their data were excluded. An olfactory ability score for each individual was derived from the sum of consistent responses per odor per person, Olfactory ability ranks are the same as above:

- Poor (0): severely impaired or no sense of smell
- Indeterminate (1 to 3): further testing required
- Normal (4-5 correct): normal or slightly reduced sense of smell

Differences between males and females were analyzed using the Mann-Whitney U test via *wilcox.test* in base R statistics.

### Ethical Considerations

Field research for this project was approved by the University of Roehampton Ethics Committee, approval number LSC 17/213 (JCB, KCH). Written informed consent included no further information beyond a name (signed and printed). Data collection sheets were labelled with a unique numeric ID. The unique numeric ID was not noted on the informed consent which means no personally identifying information was linked to test data. The data collection sheet included basic demographic information (sex, age, ethnicity) and free and verbal prompt responses to each of five odors per location.

## RESULTS

### Coding

The initial coding of descriptive data by two observers (JCB, KCH) was in moderate agreement: Repeat Markets, (free response κ= 0.537, verbal prompt κ= 0.500) and Repeat Labs (free response κ= 0.418, verbal prompt κ= 0.676). After discussion of discrepancies, we agreed to code only veridical terms as correct (even if given with other terms), unrelated food types and qualitative responses as incorrect rather than far miss, onion as near miss for garlic, and /mint as correct for peppermint. After recording, coders were in perfect agreement (κ=1).

### Field study

For the field study (n=29), we used social media (e.g., Twitter, meetup.com), blogs (smellofevolution.com), word of mouth, and direct solicitation for participants. Testing was conducted on two separate days with two groups per day (10am and 2pm). We used the area by the Tate Turbine Hall as a low odor control (low pollution, no food vendors, leafy area) for comparison to Southwark Bridge pedestrian area (traffic pollution, urine, cigarettes, and some food odors) and Borough Market (raw and cooked food vendors). We expected pollution to lower scores at Southwark and the odor signal to odor noise ratio at Borough to lower scores. The logistic regression indicates no significant effect from location (Table 1). A visualization of the change in the proportional odds (e.g., probability of being poor versus normal if you are indeterminate) show the opposite trend to what was expected. The odds of being ‘poor’ and ‘indeterminate’ decrease from Tate to the other locations (1=Tate, 2=Southwark, 3=Borough) while the odds of being ‘normal’ increase (Figure 1).

**Figure 1:**
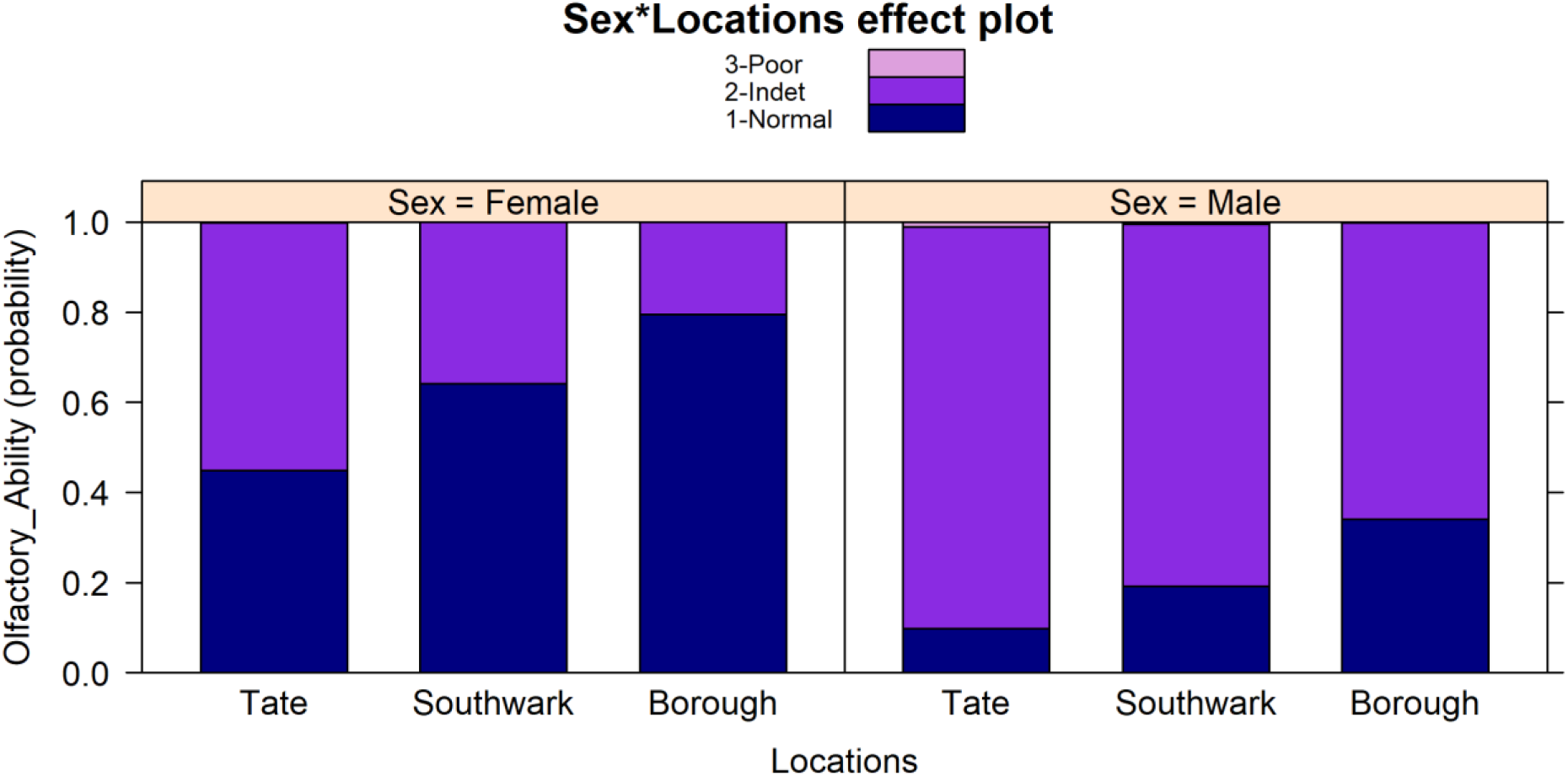
Proportional odds of changes in olfactory ability due to the environment, field study.

The only significant fixed effect is within-subjects. The difference between a model with and without the within-subjects term is significant (Likelihood Ratio statistic=16.6, df=1, p-value=0.000045). Most individuals vary from the sample average predicted response (set to 0) taking into account the estimated impact from fixed effects (Figure 2). While the significant random term in the regression indicates that individual variation impacts the model, it does not indicate if individuals vary in their scores across locations. Since there are no confounding factors from fixed effects, we used a paired ranks test to determine if there were significant changes in olfactory ability by individual (Table 2). There were no significant changes by individual for olfactory ability but there are two findings worth noting. First, the environmental impact on male olfactory ability is variable with some improving and some declining. So, while the summary statistics suggest no change, within-subjects change is taking place. Second, the environmental impact on female olfactory ability is consistent—all females improve at Borough (a dynamic food odor environment).

**Figure 2:**
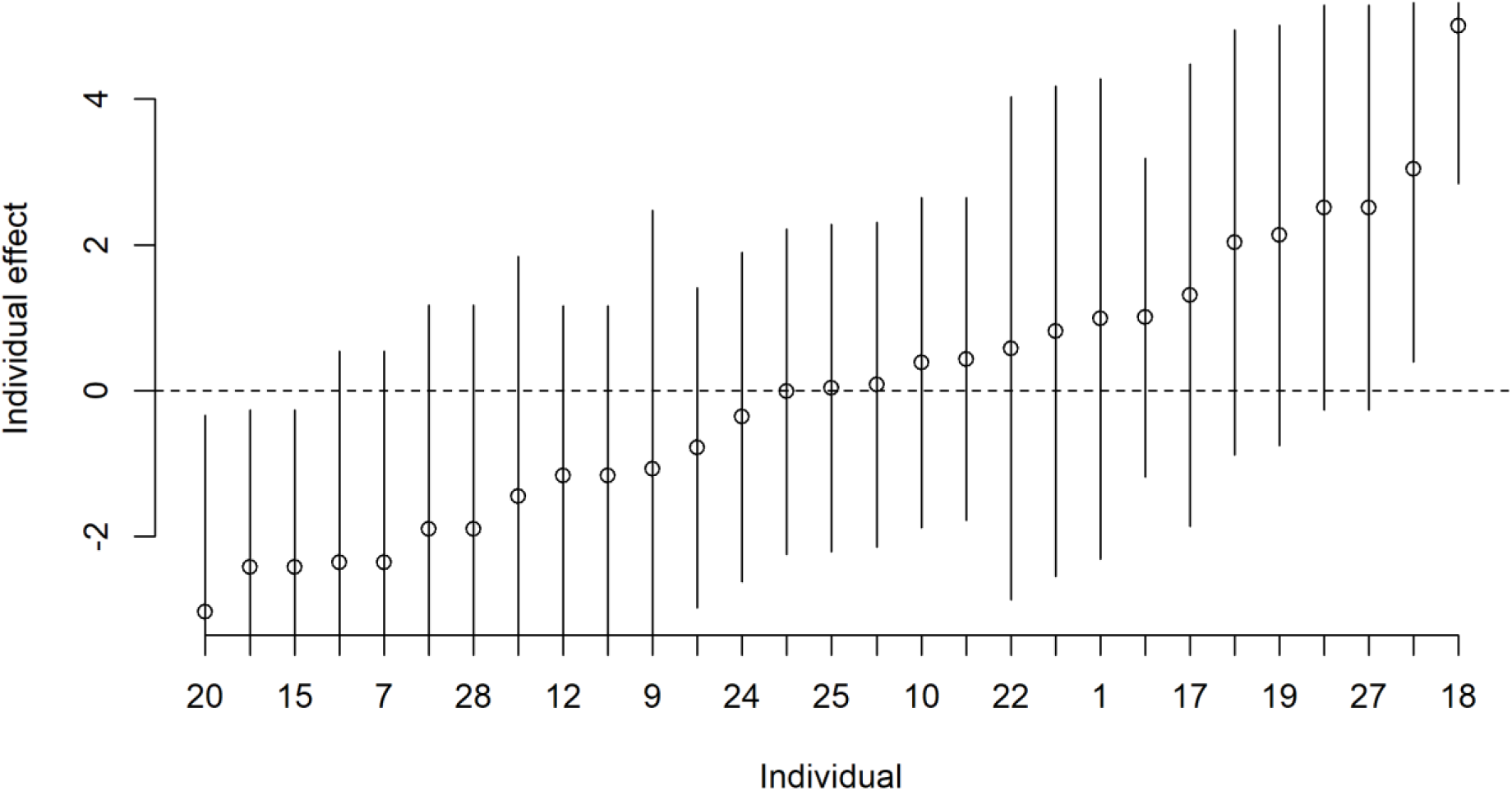
Individual effect on olfactory ability (conditional modes with 95% confidence intervals based on individual variance across locations), field study.

### Lab Study

For Repeat Labs (n=30), we recruited University of Roehampton students and staff to participate in a lab-based experiment. Testing was conducted in two labs at the same time each day over three days. Olfactory ability in a control lab (well-ventilated low-odor) were compared to those taken in an experimental condition (food market odors). Odors (garlic, coffee, madras curry paste, and caramel furanone [3% concentration]) were diffused each day one hour prior to testing. We expected olfactory ability to be negatively impacted by the experimental condition but this was not the case (Table 1). The odds of dropping in olfactory ability rank under experimental conditions (Figure 3) is slight and not significant.

**Figure 3:**
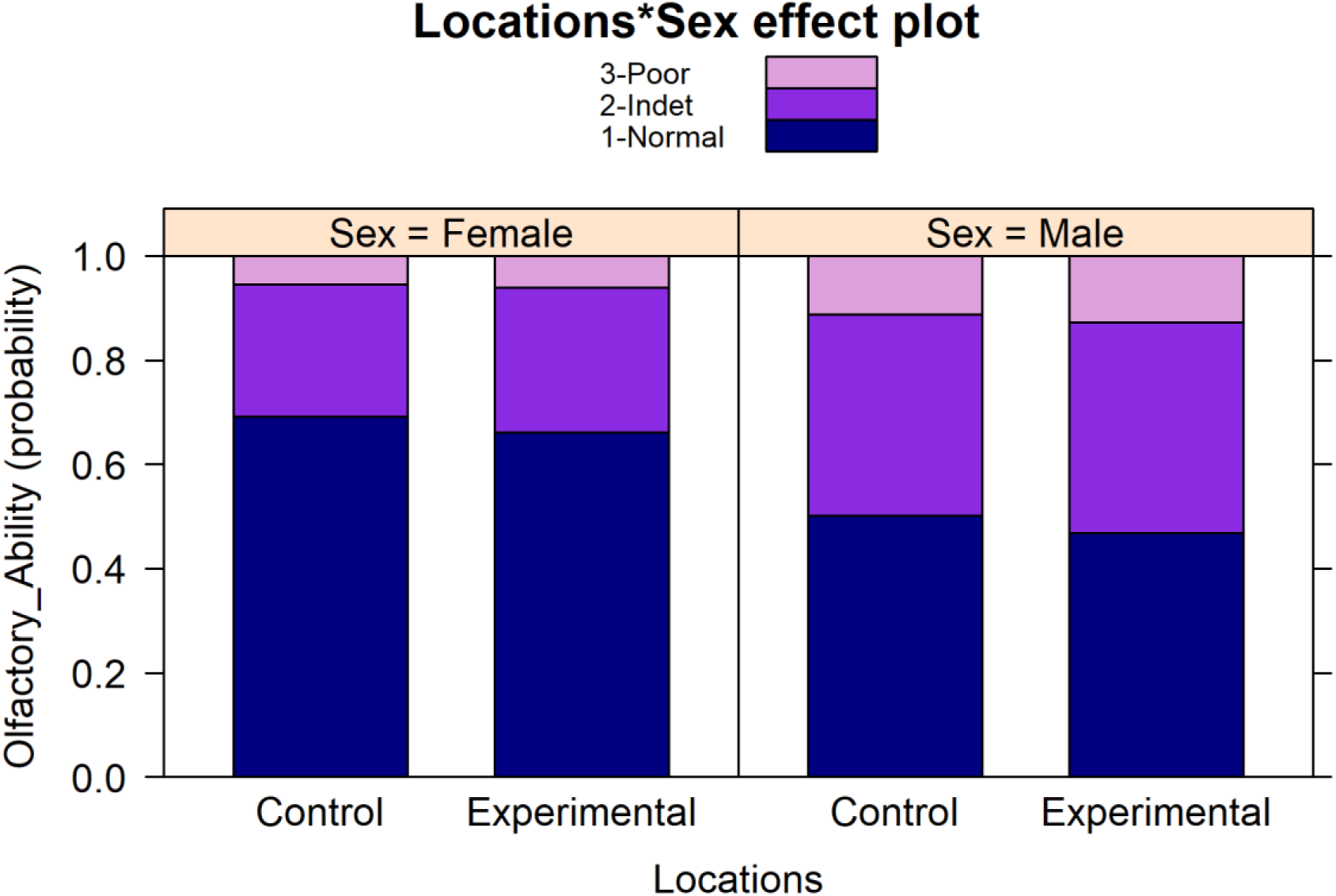
Proportional odds of changes in olfactory ability due to the environment, lab study.

Again, the only significant model term is within-subjects. The difference between a model with and without the within-subjects term is significant (Likelihood Ratio statistic=3.74, df=1, p-value=0.053. All individuals differ from the average predicted response to varying degrees (Figure 4). The paired ranks test did not return a significant result but the direction of change is variable across the sexes—while females tend to improve more than decline, the positive directionality of the field study is not replicated.

**Figure 4:**
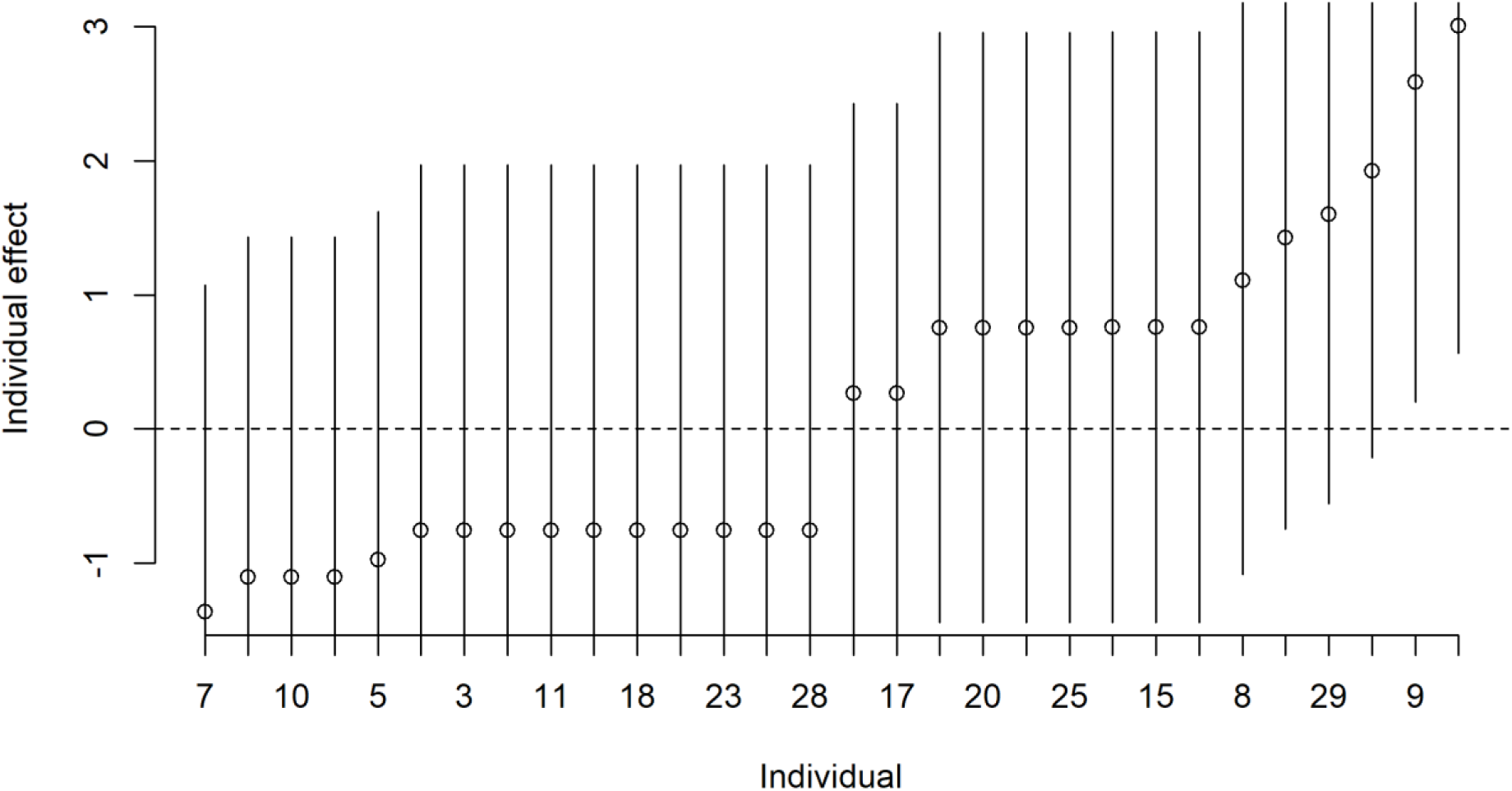
Individual effect on olfactory ability (conditional modes with 95% confidence intervals based on individual variance across locations), lab study.

### Field and Lab Combined

Despite the lack of significant differences among locations, the overall trend across both studies is that the control lab provided the best environment for olfactory ability (Figure 5). In addition, females consistently outperformed males (Figure 5) with median and modal scores of 3 (normal) compared to male median and modal scores of 2 (indeterminate). Females captured a proportionately larger share of the normosmic diagnosis: 38% of results across trials were female and normosmic but only 14% were male and normosmic. While the field settings have a much lower probability of normosmia, females have a higher probability of normosmia and a very low probability of anosmia, compared to males.

**Figure 5:**
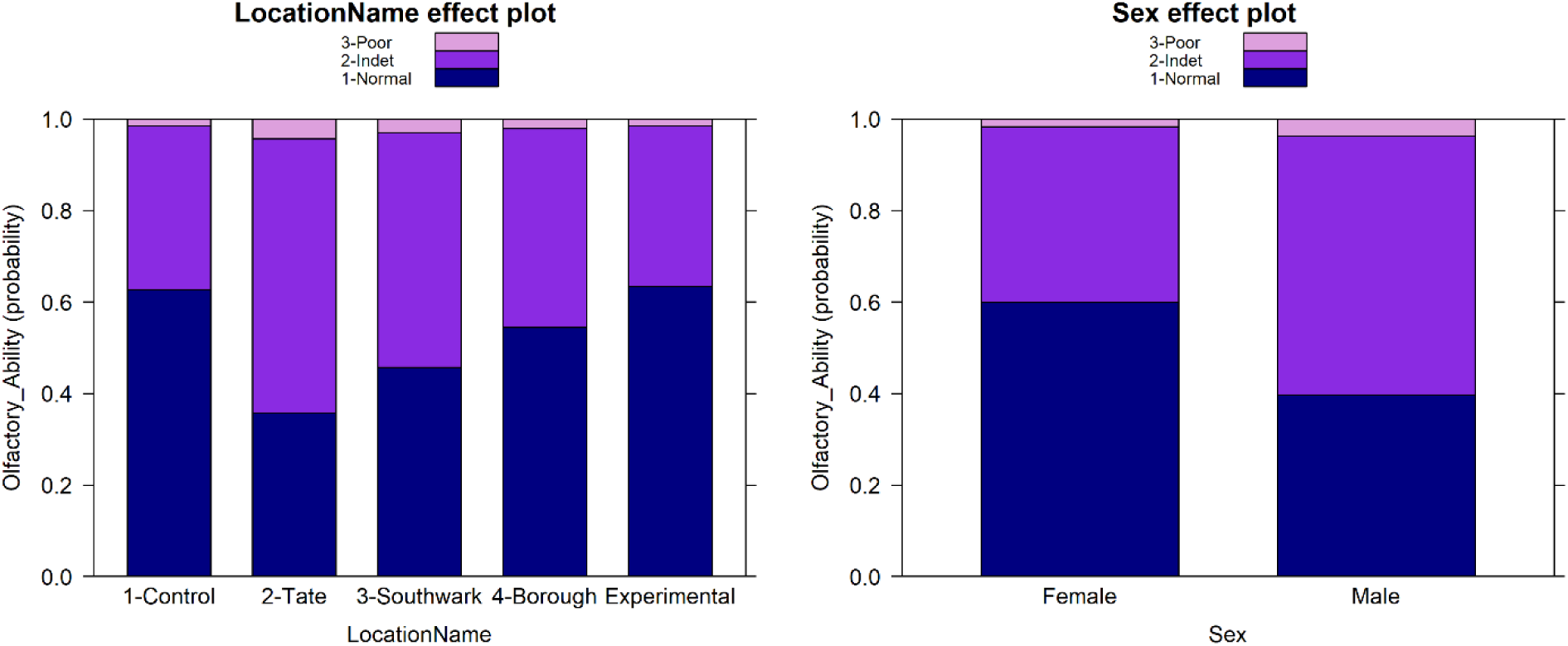
Proportional odds of changes in olfactory ability due to environment and sex.

### Consistency

There were no significant differences in olfactory ability when using the consistency scores in either study: Field (W = 103, p-value = 0.7, CI: -0.999-2.00) or Lab (W = 169, p-value = 1, CI: - 1-1). In other words, males and females are not significantly different in olfactory ability scores when responses are rated based on consistency in use of terms (e.g., chewing gum for orange across all trials), rather than correct identification of odor using veridical terms (e.g., orange for orange). Some odors were more consistently described than others (Figure 6). Orange was the least consistently described odor for both males and females in Field and Lab studies (but males in the lab had an equally difficult time describing garlic consistently). Fish was the most consistently described odor for females in the Field and peppermint in the Lab. Garlic was the most consistently described odor for males in the Field (with peppermint a close second) and fish and garlic the most consistently described odor for males in the Lab.

**Figure 6:**
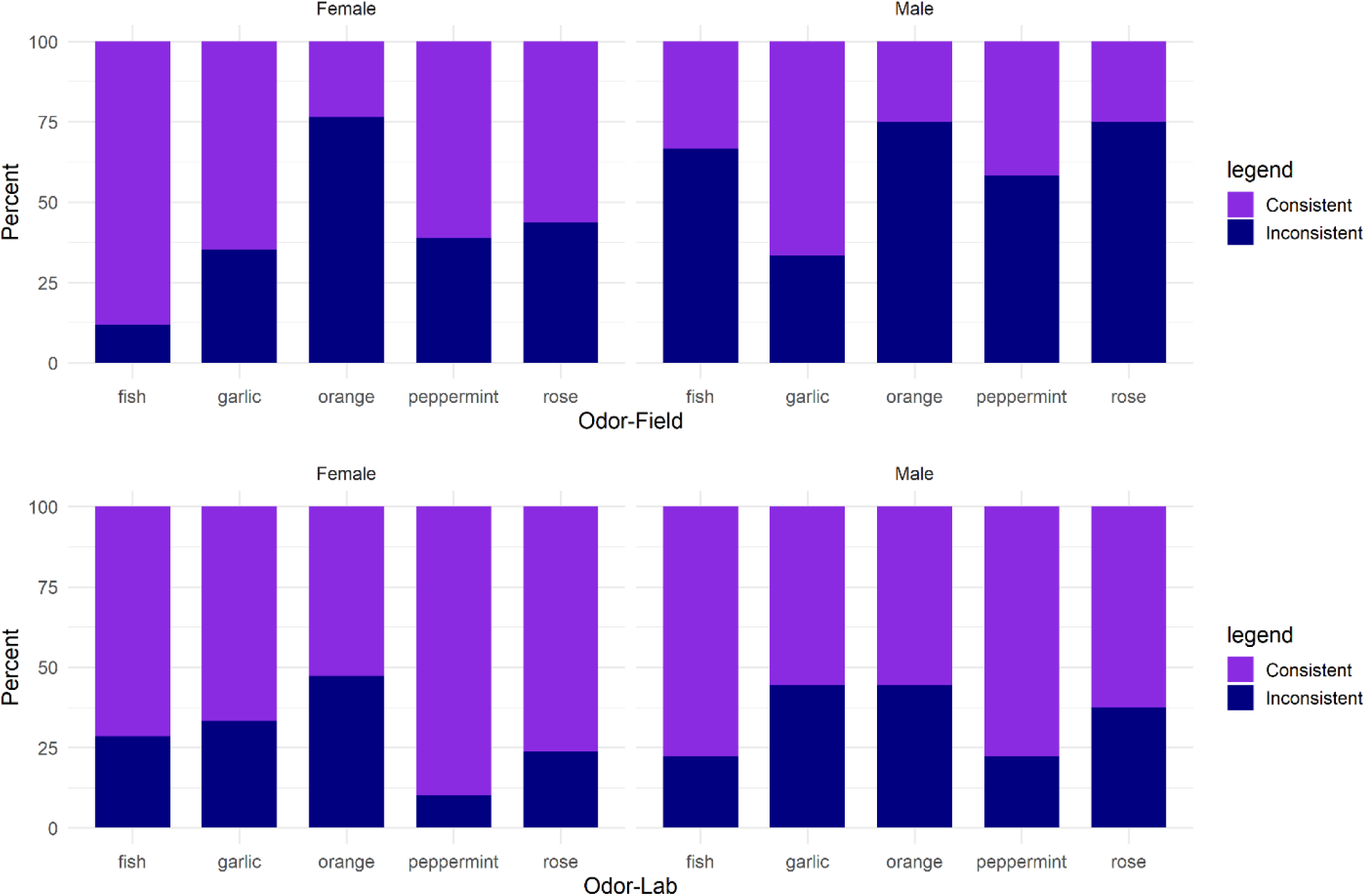
Variation in odor label consistency by sex.

For the Field study and individuals who used a consistent term two of three times, we examined which of the three locations had the most inconsistent terms (e.g., consistency at Tate and Southwark but not Borough). Females were most inconsistent terms at Tate (n=18, 66% of the time) compared to Southwark (n=4, 15% of the time) and Borough (n=5, 19% of the time). Males exhibited no pattern: Tate (n=0, 39% of the time), Southwark (n=7, 30.5% of the time), Borough (n=7, 30.5% of the time).

## DISCUSSION

Our main goal was to conduct preliminary data collection on how humans smell in the dynamic odorscapes of everyday living—how well are they able to recognize ecological salient odors (in our study, food) in a variety of settings. The difference between lab-based olfactory ability data and the limited dataset collected in the field indicates a large effect size introduced by the non-lab environment on odor sensitivity and discrimination test scores (Sorokowska et al., 2015; Sorokowska et al., 2014; Sorokowska et al., 2013). Even though we were using odor identification, those data provided a target sample size for our study, which also different in one other important methodological issue—we used a repeated measures design and the other study did not. We failed to find an effect from the environment on odor identification ability—while the comparison of results across three settings failed to find a significant effect from environment on olfactory ability, the data visualization suggested it was better in the lab than the experimental setting or the three outdoor locations. Given the previously discussed problems with odor identification, perhaps our sample was too small for this particular test. We would tentatively argue that comparing lab-based data to field-based data is complicated.

Because there is no tool for mobile rapid collection of olfactory ability, our other goal was to test the use of a clinically validated five-item odor identification test—even if only to provide a rough measure of olfactory ability. Previously identified methodological problems with odor identification that affected the outcomes. First, we found that participants were learning—they relied on verbal prompt terms after the first test, even when incorrect (choosing raspberry for orange consistently) and ignored the card in subsequent tests (‘same as before but stronger’, ‘same as last time’). Perhaps longer between-test gaps would mitigate the effect but would require greater participant commitment to the process. Second, observer coding of odor terms introduces subjectivity to the process of marking answers as correct, near miss, far miss, or incorrect (Kern et al., 2014). Our first test for inter-observer agreement suggested the two coders were in moderate agreement on marking. Our discussion revealed where the key differences lied and, once we agree on marking terms, we were in agreement. So, while there was negotiation to come to agreement, there was subjectivity that makes replication challenging in any assessment of odor identification (Rabin et al., 1984). Data on social consensus for odor terms and usage frequency of those terms would help mitigate marking correct choices as incorrect—such as knowing that a population associates the eugenol with the dentist, rose or lemon with cleaning supplies, or peppermint and orange with chewing gum—but those data are not routinely collected. Third, using an alternate approach to code ability based on the consistent use of odor term labels across trials (Rabin et al., 1984) resulted in a different outcome than the primary method of assessment using veridical terms—the sex-based differences in olfactory ability were not found when considering consistency. The difference in results is easily explained by clinical data that suggest females are much better at odor identification tasks (Richard L. Doty, 2001)— this is true even when males and females are asked to remember odor terms in a foreign language (Dempsey et al., 2002). Namely, the female advantage in naming is removed when consistency is the primary rubric. In our study, females who are not consistent across three trials but are consistent across two trials tend to be consistent after first exposure to the verbal prompt—they have learned the answer.

Do our results and interpretation suggest the odor identification test is not accurate? No, the test is more likely providing information about a different cognitive process. The cerebellar subregion involved in higher-order cognitive processing for language and memory—key areas for odor labelling (i.e. language)—is associated with olfactory identification but not detection or discrimination (the more reliable tests of olfactory ability) (Wabnegger et al., 2019). The differences we found when focusing on correct source identification as compared to consistency in recognizing the same odor (whether correct or not) indicate the importance of the analytical approach relative to the research questions. We are interested in functional olfactory ability, not language or symbolism or consistency in labelling an odor. The difference is simple: being consistent in using an odor label or having an odor rich cosmology does not indicate a functional olfactory ability but naming the source is. For example, consistently labelling mercaptan (the odor added to natural gas) as eggy or smoky three times in a row will not result in a functional behavioral output in the presence of that smell in the environment, namely to take evasive action. The correct identification of mercaptan as the smell of natural gas will lead to functional behavioral output. Thus, analysis of odor identification labels that focus on correct answers (naming the source) assesses ability to identify and response to environmental information that is functionally important.

Clinical studies on western societies tend to explain sex-based differences in olfactory ability as derived from either gender-based labor or sex-based biological differences (Dempsey et al., 2002). There certainly are sex-based differences in biological structures. First, females have roughly double the number of olfactory bulb cells, both glial cells and neurons (Oliveira-Pinto et al., 2014). Second, females have a stronger olfactory modality than men (Pamela Dalton et al., 2002; Richard L. Doty et al., 1985; R. L. Doty et al., 2009; R. L. Doty et al., 1984; Oliveira-Pinto et al., 2014), even when accounting for different cultural settings in industrialized societies (Richard L. Doty et al., 1985) and this does not appear to be based on hormonal differences because male-to-female transsexuals taking gender-affirmation hormones do not exhibit the same pattern (Kranz et al., 2019). There are also differences in experience that may act on female olfactory ability. Women are more interested than men in smell (Seo et al., 2010), are better than men at learning odors (R. Doty et al., 1984; Ferdenzi et al., 2013; Larsson, 1997; Larsson et al., 2003; Lehrner, 1993), and more proficient than men at odor identification tasks (Oberg et al., 2002). The most interesting question that emerges is whether there might be an evolutionary explanation for sex-based differences in olfactory ability.

A gendered division of labor describes most of human prehistory and, even if in a modified form, contemporary societies. So, by focusing specifically on differences between men and women in food acquisition in daily tasks, we may gain some insights into either behavioral or biological selective pressures acting on female olfactory ability. Turning to our closest relatives, female chimpanzees forage alone but in spaces that overlap with other females, which generates some competition for space and food resources. High-ranking females tend to have preferential access to resource-rich sites which correlates with greater reproductive success (Pusey et al., 2013). So, reproduction is, in part, determined by foraging ability in chimpanzees. Females also more frequently use tools for food foraging and processing, a trait shared with our closest relatives, chimps and bonobos (Gruber et al., 2010; Pruetz et al., 2015), which suggests selective pressure from activity predates the *Homo*-*Pan* split. Studies of hunting ability in indigenous hunters that have included a sensory component (vision and audition) indicate that neither plays a strong role but strength does, especially upper body strength (Apicella, 2014; Stibbard-Hawkes et al., 2018). So, despite a few ethnographic studies that have focused on the rich odor symbolism of some hunter-gatherer societies (for an overview, see Classsen et al., 1994) and a study in Malay hunter-gatherers that focused on the increased prevalence of abstract odor terms in hunter-gatherers (Majid et al., 2018a), there is no information on the functional role of olfaction in hunting, let alone other activities. Further the significance of odor terms is complex—despite the increased use of abstract odor terms, smell is the least codable sense in the Malay and, compared to other populations, olfaction is relatively low in their sensory hierarchy (Majid et al., 2018b). There is, however, information on the functional role of olfaction in foraging activities. Women outperform men at foraging tasks that involve spatial memory and navigation skills (McBurney et al., 1997; New et al., 2007) and the spatial encoding of gatherable foods into memory is chronically active in women but only active under explicit motivation in men (Krasnow et al., 2011). An ability to map food resources and navigate a catchment area is adaptively advantageous. Finally, as noted above, females have a greater density of olfactory neurons than males—when coupled with the fact human olfactory sensory neurons are evolutionarily biased towards food-related odors, superior olfactory ability may well be linked to food foraging (Saraiva et al., 2019). The main takeaway is that females appear to have better signal-to-noise ability when detecting, identifying, and discriminating among food odors, a skill that provides an adaptive advantage in successful foraging and group provising—such as seen in our data when female olfactory ability was enhanced in the rich food-odor environment of Borough Market.

In conclusion and despite published data indicating a significant environmental effect on olfactory ability (Sorokowska et al., 2014; Sorokowska et al., 2013), we failed to detect any significant differences in two repeated measures studies. We suspect that either within-population repeated measures tests have a more subtle impact from environment on olfactory ability than cross-cultural comparisons or that the comparison of field- to lab-collected data overemphasized differences. We also found that odor identification is a less than ideal tool for assessing olfactory ability despite ease of delivery and speedy test time. We argue that field-based methods are needed to characterize representative variation in human olfactory ability and the importance of ecological context. Lastly, we find that even in the field and despite methodological problems, females exhibit better olfactory ability and that may be an artifact of long-standing evolutionary pressure acting on a mosaic of traits related to food foraging. This last point is important and highlights the need for field-collected data (i.e., non-lab settings), which may seem superfluous and lacking utility to those accustomed to lab environments. The reason is straight forward: the clinically-derived dataset is insufficient to capture variation in and the breadth and context of human olfaction. Indeed, compared to the traditional lab environment, a virtual reality lab environment found that behavioral responses to odors is context dependent— in the absence of ecological validity, motivated behavior is not observed (de Groot et al., 2020). Field-based data collection will widen participation beyond current olfactory ability datasets that are comprised of non-representative samples. Why non-representative? First, WEIRD samples are biased by individuals who can afford the costs of travel and time to be tested in a lab. Second, WEIRD populations (or even simply EIRD) are a global minority. Third, many populations are geographically isolated from lab-testing sites—this describes the majority of indigenous, traditionally living, small-scale societies. Fourth, lab conditions cannot be recreated in the field. Clinical data serve their purposes but field data are needed to address an entirely different question: are meaningful biological differences in olfactory ability explained by ecological context? This study is one of a small number that use contemporary validated methods of assessing olfactory ability in non-lab settings. We know very little about the ecological or evolutionary pressures acting on human olfactory ability and if we are to truly understand the extent of and variation in olfactory ability, we need better field methods and cross-cultural data as well as transdisciplinary efforts.

## ACKNOWLEDGEMENTS

Roehampton University staff and students for their support.

## CONFLICT OF INTERESTS

There are no conflicts of interest.

## AUTHOR CONTRIBUTIONS

KCH and JCB conceived and designed all studies. KCH, JCB, DB, PF conducted field-based data collection. KCH, JCB, SA conducted lab-based data collection. KCH analyzed data and drafted the manuscript. JCB revised the manuscript critically for analytical approaches and intellectual content.

## Notes

#### Summary of Updates

This version is the final submission that includes new figures done using a color blind palette that is consistent across all figures.

